# Lethal thermal limits in embryos of two intertidally brooding nassariid species, and evolution of a protective developmental mechanism conditioned by maternal experience

**DOI:** 10.1101/2023.05.15.540845

**Authors:** Morgan Q Goulding

## Abstract

*Tritia obsoleta* and *Phrontis vibex* are distant relatives in the neogastropod clade Nassariidae (1). Both species are common along coastline from Maine to the Caribbean; females deposit clutches of zygotes in capsules cemented to solid surfaces in intertidal flats (2), with swimming veliger larvae emerging after about a week of development. In this study, embryos of *T. obsoleta* (from Massachusetts and Florida) and *P. vibex* (from Florida) were exposed to temperatures ranging from 33° to 37°C. In both species, very young embryos are especially sensitive to thermal stress. Brief early heat shock did not disturb spiral cleavage geometry but led to variable, typically severe defects in larval morphogenesis and tissue differentiation. In *T. obsoleta* but not *P. vibex*, early heat shock resulted in immediate slowing or arrest of interphase progression. This reversible arrest is correlated with improved prognosis for larval development, and (in Massachusetts snails) depends on parental exposure to warm temperature. Embryos from Massachusetts snails housed at lower temperature exhibited cytokinesis defects in response to even mild warming. Thus, parents informed by experience may or may not endow their young with a developmental braking mechanism that lets them withstand occasional perilous conditions during the most vulnerable stage of life.

## Results and Discussion

*Phrontis vibex* and *Tritia* (formerly *Ilyanassa*) *obsoleta* are winkles with shared habitats and similar life histories. Basally related within the clade of Nassariidae, their lineages are estimated to have diverged roughly 58 million years ago (1). The easily accessible embryos of the two species show practically indistinguishable complex spatiotemporal patterns of morphogenesis from egg meiosis through larval hatching (MQG, unpublished). Embryos in this study were provided by snails from three sources. *P. vibex* was collected on Mashes Sands Beach (29.971, -84.352) in the Gulf of Mexico. *T. obsoleta* was collected from Amelia Island (30.628, -81.484) in the Intracoastal Waterway, and another group of *T. obsoleta* was obtained from the Marine Biological Laboratories, Woods Hole, Massachusetts. Snails were kept in aerated artificial seawater (Instant Ocean, ∼25/gallon) sufficient to nearly fill 6-quart shoeboxes with about an inch of sand, in a warm room (25.5°C ±1°C). A subset of the Massachusetts *T. obsoleta* was kept exclusively in a cool room (16°C ±1°C). Embryos were cultured at 25°C, except where otherwise stated.

In a preliminary experiment to test the upper limit of temperature compatible with development, 34 intact *P. vibex* egg capsules were incubated for 84 hours at 37°C, beginning 6-24 hours after capsule deposition. At the end of this time, sixteen capsules (with 20-30 eggs/capsule) were full of nearly hatching-stage veligers with no detectable abnormalities; in the other eighteen capsules, every single embryo completely failed to differentiate any tissues. This stark result implies that mammalian body temperature ruins the development of *P. vibex* in a stage-specific manner, with the very young (less than 12-18 h post-oviposition) being extremely vulnerable to heat shock. In another set of *P. vibex* embryos removed from their capsules at the one-cell stage and cultured seven hours at 37°C, no abnormal cell morphology was observed at the 12-to 16-cell stage or the 24-to 29-cell stage (n=63). These results suggest that some heat-sensitive factor(s) might be involved in early steps of cell fate determination.

Effects of early heat shock were examined more carefully in another experiment using decapsulated clutches of embryos, with Florida *T. obsoleta* embryos treated in parallel to *P. vibex*. A six-hour incubation at 37°C starting at the 2-cell stage yielded highly variable, generally severe defects in larval morphogenesis and differentiation in 92% of *P. vibex* embryos (n=35), but only in 67% of *T. obsoleta* embryos (n=47). This temperature, if experienced during very early embryogenesis, thus appears to approximate the upper limit for life in the latitudinally equivalent populations of both species. In further experiments with Florida *T. obsoleta*, early (1-or 2-cell) embryos incubated 4 h at 35°C developed normally to hatching stage (>90% of the eggs from six capsules).

During these experiments I noticed that cell divisions seemed to stop during high temperature incubation in *T. obsoleta*. Immediately following a six-hour incubation at 37°C, begun during interphase of the 2-cell stage, 100% of *T. obsoleta* embryos were arrested as 4-cells (n=39), while all *P. vibex* embryos had reached the stage of 24-29-cells (the same stage as sibling controls kept at 25°C) (n=13). Further, I found that *T. obsoleta* early embryos incubated at 35°C do not arrest, but significantly slow their cell cycles to over twice the normal period. Time lapse imaging at 35°C showed the length of cell division delay to be highly variable within a clutch, with a range of over 40 minutes. This retardation appears to reflect a delay of M phase onset: while the total cell cycle period (first cleavage to second cleavage) was increased by ∼110% to 150%, M phase duration (from the time of polar lobe furrow ingression to relaxation) was not significantly altered.

Unsurprisingly, *T. obsoleta* from Massachusetts is much more heat-sensitive than *T. obsoleta* from Florida. However, embryo heat-tolerance appears to depend strongly on parental snails’ temperature-adaptation. These facts were revealed first by an experiment in which gastrula-stage *T. obsoleta* embryos from three sources were cultured at 33°C for three days: those from either Florida or Massachusetts adults kept in the warm room (1 clutch of eggs from each) developed normally to fetal veliger stage (100%); by contrast, embryos from Massachusetts adults kept in the cool room (four clutches) invariably developed as monsters with multiple organogenesis defects. In another experiment, embryos from the same three sources were cultured from early development (1-to 2-cell stage) at 33°C for three days. Development of Florida embryos was 100% normal (6 clutches). Embryos from warm-adapted Massachusetts parents permanently arrested at very early cleavage stages (100%, 5 clutches). Embryos from cool-adapted Massachusetts parents developed as variably malformed monsters with stunted, misplaced, and missing organs (100%, 5 clutches). Hence it appears that unusually warm ambient temperatures (25°C, unusually warm for Massachusetts) cue *T. obsoleta* parents to equip their embryos with an arrest mechanism that protects against thermal stress. This mechanism is clearly adaptive only in a situation of transient warming, as would normally be experienced at low tide.

In another experiment, three clutches of eggs from cool-adapted Massachusetts *T. obsoleta* were heat-shocked 5 h at 35°C, starting at the 1-or 2-cell stage. In two clutches, all embryos exhibited reversible retardation; in the third clutch, however, mitosis began on schedule, and nearly every embryo failed to complete cytokinesis. A penetrant cytokinesis defect was also witnessed in early (1-cell) embryos from another clutch of cool-adapted Massachusetts *T. obsoleta* during incubation at only 33°C. From these results it can be concluded that (1) cool-adapted *T. obsoleta* are not absolutely incapable of endowing their eggs with a protective arrest mechanism, but seem less likely to do so, compared to warm-adapted parents; (2) reversible retardation protects early embryos from death by cytokinesis failure as well as death by cell fate specification defects.

Clearly, brief exposure to temperatures above a population-specific normal environmental range disrupt multiple developmental processes in nassariid embryos. *T. obsoleta* parents respond to a warm environment with an arrest mechanism that protects embryos at the most heat-sensitive stage (i.e., cleavage). The nature of this arrest (prolonged interphase) seems well-suited to the physiological threats (cytokinesis failure, or mis-specification of regional founder cells during cleavage). Comparing results of brief vs. prolonged incubation at 33°C suggests that the duration of cell cycle arrest may be limited, depending on the difference between temperatures experienced by the embryo and the parent(s). Pronounced variability in heat stress resistance (including the mitotic delay mechanism) was evident between clutches, suggesting that some individuals are better prepared than others to protect offspring from new extremes of warmth. The protective mitotic delay discovered in *T. obsoleta* may represent an evolutionary novelty within the Nassariidae, possibly associated with a transition to living in the intertidal zone (1).

This work was supported by the Thomas Biology Chair Fund of the GSW Foundation at Georgia Southwestern State University.

